# The mitoribosome-associated factor Mrx9 acts as a negative regulator of the prohibitin/m-AAA complex

**DOI:** 10.64898/2026.07.28.741247

**Authors:** Jhulia A. C. Chagas, Flavia Fontanesi, Amanda C. Pereira-Antonio, Mário H. Barros

## Abstract

The synthesis of mitochondrial-encoded polypeptides is an essential process, primarily regulated at the posttranscriptional level. In yeast, many regulatory factors have been described as acting in association with the mitoribosome to promote efficient translation; however, the precise mechanisms by which these components function remain largely unknown. Here, we expand on findings concerning a previously studied mitoribosome interactor, Mrx9, which is found in large expressosome-like assemblies of mitoribosome clusters. Mrx9 was initially linked to mitochondrial translation and was suggested to be necessary for splicing of *COX1* and *COB* transcripts. Previous data obtained by others in proximity labelling assays demonstrating that Mrx9 is in close proximity to both the mitoribosome exit tunnel and the prohibitin (PHB) complex. Our current experimental data show that overexpression of Mrx9 impairs the proteolytic functions of Yta10 and Yta12 within the prohibitin complex, leading to splicing defects, accumulation of aberrant polypeptides, and a noticeable impairment in the processing of the essential mitoribosomal protein bL32m. These findings support a regulatory role for Mrx9 in the PHB/m-AAA complex by modulating the activities of both Yta10 and Yta12.

## Introduction

Mitochondria contain their own set of ribosomes (mitoribosomes), which are highly specialized in the translation of hydrophobic polypeptides, the core subunits of the oxidative phosphorylation system (OXPHOS). Despite sharing certain features with bacterial ribosomes, mitoribosomes differ markedly from other ribosomes in both structure and composition [1,2]. In *Saccharomyces cerevisiae*, mitoribosomes are spatially organized into higher-order assemblies known as MIOREX (*<u>Mi</u>tochondrial <u>Or</u>ganization of Gene <u>Ex</u>pression*) [3]. This sophisticated arrangement forms large complexes that integrate a wide range of factors tightly connected to gene expression events, such as RNA splicing and processing, translational activation, mitoribosome biogenesis, and RNA decay [3,4]. Most of these mitoribosome-associated factors have been studied over the years [5–8]; however, a set of them remain poorly understood. We have made efforts to characterize one of these unknown proteins, named Mrx9 (MIOREX component 9), which initially caught our attention due to its role in mitochondrial RNA processing and translation [9].

Previously, we showed that the loss of Mrx9 does not affect yeast growth on non-fermentable carbon sources or mitochondrial translation [9]. Conversely, overexpression of Mrx9 is linked to toxic phenotypes. An excess of Mrx9 results in low levels of *COX1* and *COB* mature transcripts as shown by northern blot analyses, and consequently, impaired translation of these polypeptides. Additionally, the toxic effects of *MRX9* overexpression depend on strains harboring mitochondrial DNA (mtDNA) introns, suggesting that Mrx9 might affect the splicing of *COX1* and *COB* mRNAs. Based on our early screenings, Mrx9 was proposed to exert a negative role on RNA processing and mitochondrial protein synthesis, because its effects were primarily observed under overexpression conditions [9]. However, the molecular mechanism by which Mrx9 functions remains unclear and needs to be addressed.

Splicing of mitochondrial primary transcripts is facilitated by protein factors known as maturases, which are encoded by reading frames located within some of the introns in the *COX1*, *COB*, and 21S RNAs [10,11]. In addition to maturases, nuclear-encoded products are required for proper mitochondrial intron splicing. Interestingly, most of these products have multiple functions in mitochondrial RNA metabolism. For example, Nam2/Msl1, the mitochondrial leucyl-tRNA synthetase, is required for the excision of the aI4 and bI4 introns of *COX1* and *COB* [12]; Pet54 was initially identified as a translational factor for *COX3* mRNA [13], but was later shown to participate in *COX1* aI5β intron excision [14]; the RNA helicase Mss116 is required for the splicing of all mitochondrial introns and mitoribosome assembly [15,16], and the mitoribosome protein Cox24/mS38 was implicated in the processing of *COX1* aI2 and aI3 introns [17].

The multiple functions of factors involved in mitochondrial RNA metabolism allow the layered regulation of mitochondrial gene expression [8,18–20]. Moreover, a key class of proteins with essential roles is represented by translational activators, which are critical for mRNA stabilization and translation initiation. Notably, many of these activators play more than one function in the complex regulatory network that controls mitochondrial gene expression, often acting within feedback control loops [21–23]. For example, translation of *COX1* and *COB* transcripts relies on a diversity of activators that operate at distinct regulatory steps, ranging from binding to the 5’ untranslated region (UTR) of mRNAs to associating with the newly synthesized polypeptides in intermediate stages of respiratory complex assembly [18,24,25]. Additionally, negative repressors are also required for refining the control of mitochondrial translation rates [6,24], not only to achieve a precise stoichiometry of OXPHOS components encoded by two different genomes, but also to avoid the accumulation of unassembled subunits with toxic redox properties [26].

In this study, we present new data showing Mrx9 involvement in mitochondrial translation. Specifically, we observed that a tagged version of Mrx9 pulled down the prohibitin (PHB) complex and the membrane-bound-ATPases proteases (m-AAA), which are associated with diverse cellular activities [27]. The PHB/m-AAA complex is an essential quality control machinery for all newly synthesized polypeptides [28]. Our evidence, and previous proximity studies [28, 29], suggest that Mrx9 is located at the polypeptide exit tunnel (PET) of the mitoribosome, likely as an accessory component of the PHB/m-AAA complex. The PET is a site in the mitoribosome large subunit (mtLSU) that is critical for protein synthesis, where newly synthesized polypeptides emerge and undergo immediate folding and insertion into the IMM [1,30]. Here, we show that *MRX9* overexpression impairs the m-AAA proteases functions within the organelle, including intron excision, proteolysis of aberrant translation products, and partially affects the maturation of the mtLSU component bL32m, suggesting that Mrx9 might act as a negative regulator of the Yta10 and Yta12 proteases.

## Results

### Overall effects of *MRX9* overexpression on mitochondrial protein synthesis

In a previous overexpression screen, we found that an excess of Mrx9 specifically impairs Cox1 and Cob expression [9]. To confirm and expand this observation we repeated the *in vivo* metabolic labeling of mitochondrial products, comparing wild type, Δ*mrx9,* and *MRX9* overexpression under the control of the *GAL10* promoter (Figure 1A). In cells overexpressing *MRX9*, we observed a decrease in Cox1 and Cob translation, in agreement with our previous findings. However, other polypeptides also exhibited reduced synthesis in an intronless strain, most notably complex IV subunits Cox2 and Cox3, and complex V subunits Atp6 and Atp8, when compared to a normal intron-containing strain (Figure 1A-B), suggesting a general impairment in mitochondrial translation under *MRX9* overexpression, which had not been captured quantitatively in our previous work [9]. The contrast between the two strains reinforces the relevance of mtDNA introns in the context of regulating mitochondrial gene expression [31]. The absence of introns intensifies Mrx9 toxicity, causing a broader translational defect that is not observed in the intron-containing strain.

**Fig. 1.**
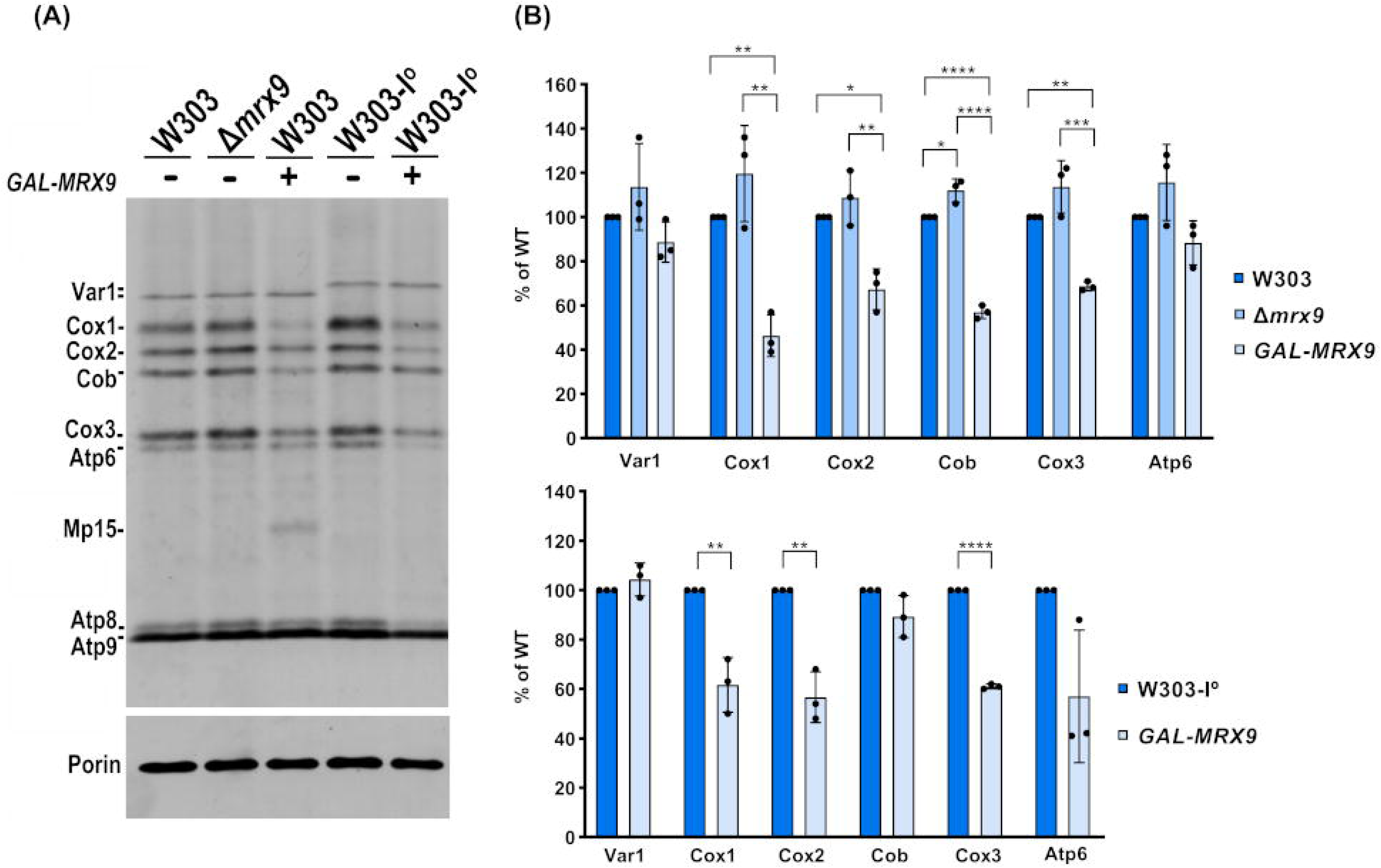
Overexpression of *MRX9* gene impairs mitochondrial translation. (A) Analysis of newly synthesized mitochondrial translation products in whole cells. The indicated strains were pulse-labeled with [³ S]-methionine in the presence of cycloheximide to inhibit cytoplasmic protein synthesis for 10 minutes at room temperature, as described in the Materials and Methods section. Porin was used as a loading control. (B) Quantification of band intensity ratios for intron-containing (W303) and intronless (W303-I) strains, normalized to porin. Tukey’s multiple comparisons test was applied for three groups and the unpaired Student’s t-test for two groups. Data represent mean ± SD (*n = 3*). *P* values are indicated as follows: *P* < 0.05 (*), *P* < 0.01 (**), *P* < 0.001 (***), *P* < 0.0001 (****).

In a strain harboring mitochondrial introns, *MRX9* overexpression results in the synthesis of Mp15, a truncated Cox1 polypeptide (Figure 1A). This truncated product is not present in cells transformed with an empty vector with the *GAL10* promoter (Figure S1A). Mp15 was first described as an aberrant product of Cox1 translation, resulting from premature termination of the *COX1* mRNA, and accumulates in various COX mutants in a *COX1* intron composition-dependent manner [32]. In our previous study, when we overexpressed 34 components of the MIOREX individually, only the excess of *MRX9* or *MRX6* resulted in a clear accumulation of Mp15 [9]. Next, we asked whether overexpression of factors involved in *COX1* and *COB* intron splicing could rescue the Mp15 accumulation phenotype caused by *MRX9* overexpression. To test this, we individually overexpressed *MNE1*, *CCM1*, *CBP1*, and *MRS1* [33—36]. In all cases except *MRS1*, Mp15 levels remained unchanged, indicating that *MRX9* excess–induced toxicity was not alleviated. In cells overexpressing *MRS1*, however, Cox1 translation was also abolished, suggesting that excess Mrs1 causes a premature defect in *COX1* expression that precedes Mp15 accumulation (Figure S1B). Overexpressing *MRX9* in an intronless (W303-I°) strain did not affect Cob translation (Figure 1A), suggesting that the observed alteration in Cob expression associated with the intron-containing mtDNA is the consequence of a splicing defect. Conversely, Cox1 translation was slightly impaired by *MRX9* overexpression, although not to the same extent detected in the intron-containing cells. In addition, the translation of Cox2 and Cox3 still appeared to be strongly affected in the W303-I° + *GAL-MRX9* background (Figure 1A-B, Figure S2). Collectively, these findings suggest the existence of two distinct mechanisms affecting mitochondrial translation upon *MRX9* overexpression. The first relates to a defective splicing, which accounts for the impaired synthesis of Cob and, in part, Cox1 polypeptides. Notably, the impairment in translation for other polypeptides is seen in both intron-containing and intronless backgrounds, pointing to a second splicing-independent mechanism of action. These data highlight that the negative features of *MRX9* overexpression are not limited to the presence of introns, emphasizing a broader regulatory role for Mrx9 in mitochondrial gene expression.

### *COX1* and *COB* mature transcripts are affected by *MRX9* overexpression

We have previously shown by northern blot analysis that mitochondria isolated from cells overexpressing *MRX9* accumulate low levels of mature *COX1* and *COB* mRNAs [9]. Because maturation of these transcripts is strictly dependent on the correct splicing of intron-containing pre-mRNA, it is important to consider that the intron composition changes depending on the strain genetic background. In the W303 yeast strain, four introns are found in the *COX1* sequence (aI1, aI2, aI3 and aI5γ); five introns in *COB* (bI1, bI2, bI3, bI4 and bI5), and only one intron is present in the 21S rRNA (Sce-I) [37]. To further investigate the effects of Mrx9 excess on the levels of mitochondrial transcripts, we comprehensively quantified mRNAs and rRNAs by RT and qPCR analyses. To detect the mature transcripts, primers were designed to amplify RNA across splice junctions so that only spliced RNAs could be amplified, while to quantify both processed and unprocessed transcripts, internal exon regions were targeted (Figure 2A). Thus, elevated levels of Mrx9 induced a clear decrease in the levels of mature *COX1* and *COB* mRNAs, especially the latter. Conversely, mature 21S rRNA showed no impairment (Figure 2B). When processed and unprocessed *COX1* and *COB* RNAs were quantified together, we observed an increase in the levels of total mRNAs compared to wild-type cells (Figure 2B), likely due to an accumulation of unprocessed transcripts. We also analyzed 15S rRNA, *ATP6* and *COX2* transcripts, which do not contain mitochondrial introns, and found no significant differences (Figure 2B).

**Fig. 2.**
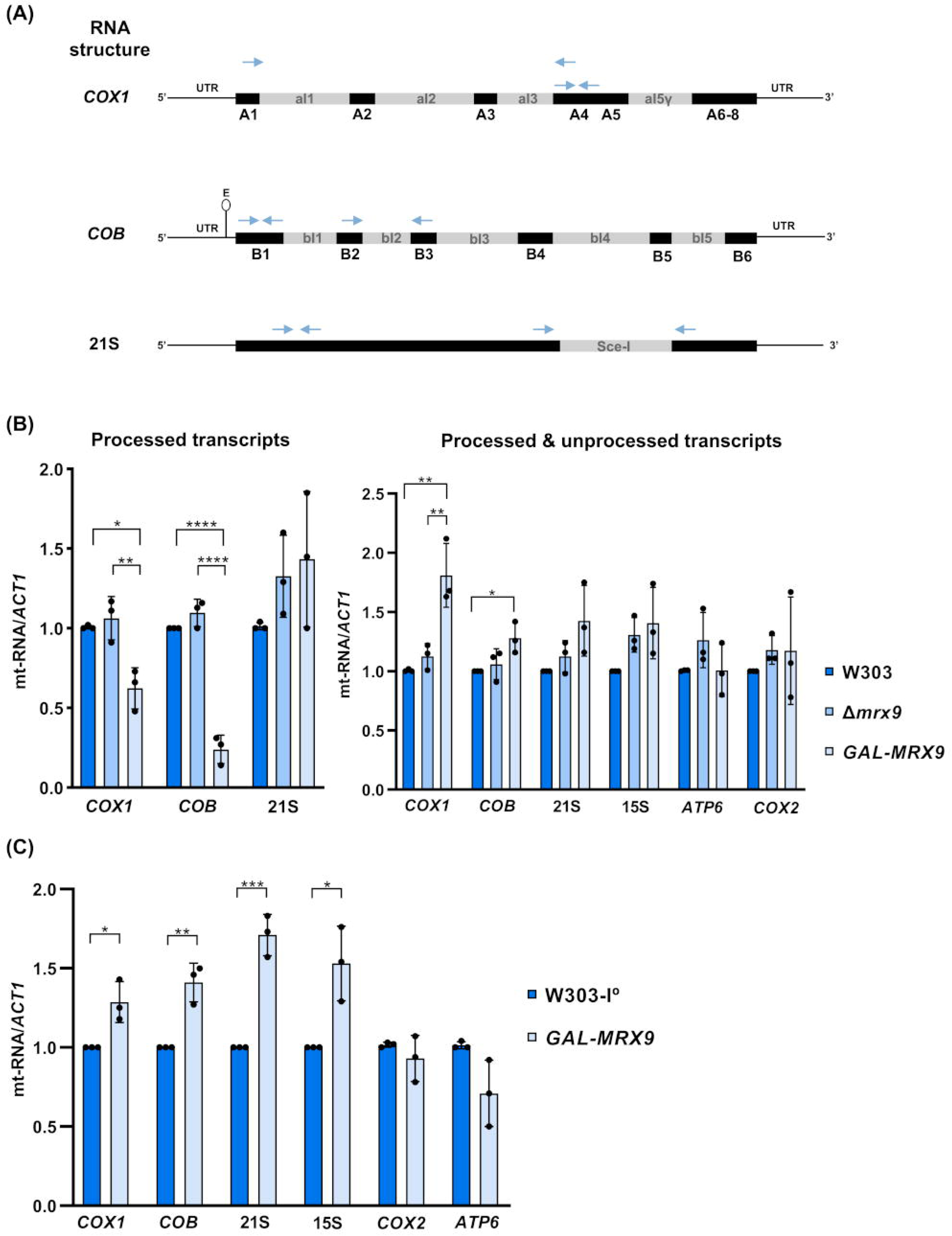
Mrx9 excess results in low levels of mature *COX1* and *COB* RNAs. (A) Schematic representation of *COX1*, *COB* and 21S rRNA transcripts in the W303 yeast strain. *COX1* (A1–A6-8) and *COB* (B1–B6) exons are shown in uppercase. The only intron in 21S rRNA is named *Sce-I*. Blue arrows indicate primer positions in intron-containing RNAs: primers spanning intron–exon junctions were designed to amplify only mature transcripts, while primers annealing within exon regions amplify both processed and unprocessed RNAs. (B) RT-qPCR analysis of mitochondrial RNA levels in WT (W303), Δ*mrx9* and *GAL-MRX9* cells, expressed as fold change relative to WT. Actin was used as the normalization control. Bars represent the mean ± SD of three independent experiments. Statistical significance was determined by using Tukey’s multiple comparisons test. (C) Same as in (B), but in strains devoid of introns in the mitochondrial DNA. Statistical significance was determined by using the unpaired Student’s t-test. *P* values are indicated as follows: *P* < 0.05 (*), *P* < 0.01 (**), *P* < 0.001 (***), *P* < 0.0001 (****).

Since *MRX9* overexpression compromised mitochondrial translation in an intronless strain (Figure 1A), we next asked whether this toxicity could be a consequence of changes in RNA availability. In the intronless background, most mitochondrial transcripts displayed higher accumulation in cells overexpressing *MRX9* compared to wild-type cells (Figure 2C), indicating that the defect in mitochondrial protein synthesis as a consequence of elevated Mrx9 levels is not dependent on mRNA or rRNA availability. Altogether, these data suggest a clear defect in the splicing of both *COX1* and *COB* transcripts in *GAL*-*MRX9* cells, while no effect is observed for the 21S rRNA processing. Remarkably, mitochondrial translation is negatively impacted by *MRX9* overexpression in intronless cells where transcript accumulation is not decreased. These results corroborate the hypothesis that Mrx9 might play additional roles in mitochondrial gene expression other than the splicing process.

### Mitoribosome biogenesis is enhanced as a consequence of high Mrx9 levels

Given the observed general effect on mitochondrial protein synthesis under high Mrx9 levels (Figure 1) and the tendency to increase rRNA levels (Figure 2), we next investigated whether these features could indicate mitoribosome subunit accumulation. Previous studies showed that increased mtLSU and mtSSU accumulation is part of a compensatory mechanism in which mitochondria attempt to respond to an inefficient mitoribosome performance by increasing biogenesis. One approach to detect such accumulation is to measure the steady-state level of representative mtLSU and mtSSU proteins [20,38]. We therefore analyzed mitochondria isolated from *MRX9*-overexpressing and wild-type cells, and tested mitoribosome proteins levels. We examined four late assembly proteins of the mtLSU (bL32m, uL16m, ul2m and uL3m), and one intermediate-stage protein (uL29m) [39]. For the mtSSU, one early assembly component (uS12m) and two late assembly proteins (uS14m and mS37) were tested [40,41]. Our results show that Mrx9 excess increased the endogenous levels of most mitoribosome proteins analyzed, indicating an increase in mitoribosome abundance (Figure 3A-B).

**Fig. 3.**
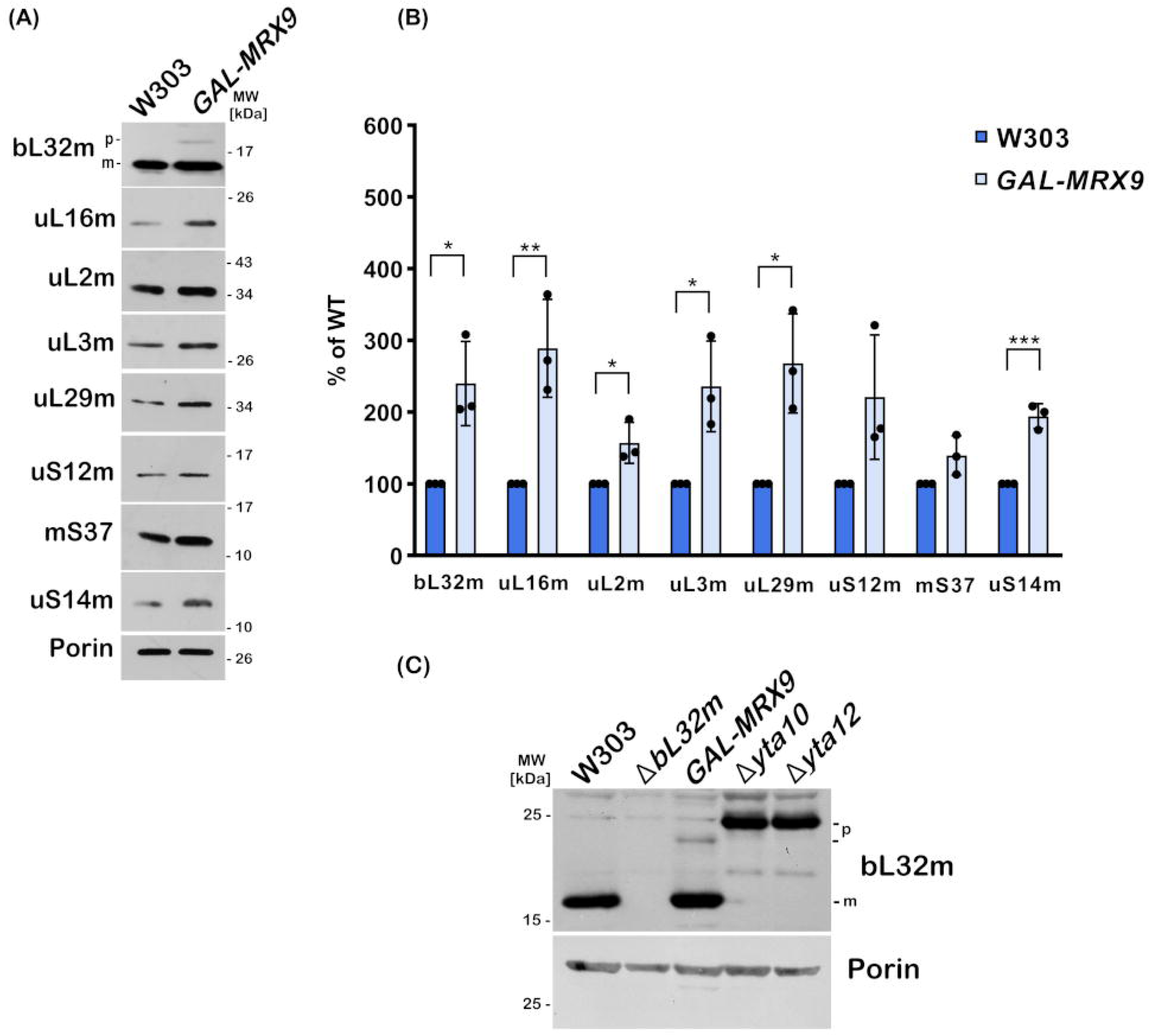
Mitoribosome biogenesis is upregulated in mitochondria isolated from *MRX9-*overexpressing cells. (A) Steady-state levels of the indicated mitoribosome proteins in mitochondria isolated from WT (W303) and *GAL-MRX9*-HA strains. 40 µg of proteins were separated by 12% SDS-PAGE. Antibodies were used against components of the LSU and SSU. Porin was used as loading control. (B) The graph represents the quantification of band intensities from the Western blot analysis. Quantification was normalized to porin. Statistical significance was determined by using the Student’s *t*-test. *P* < 0.05 (*), *P* < 0.01 (**), *P* < 0.001 (***). (C) Steady-state levels of bL32m in mitochondria isolated from WT (W303), Δ*bL32m*, *GAL-MRX9*-HA, Δ*yta10* and Δ*yta12*. 40 ug of proteins were separated by 14% SDS-PAGE. Porin was used as loading control. p, precursor bL32m; m, mature (processed) bL32m.

Among the quantified mtLSU proteins, we observed a partial accumulation of a putative unprocessed form of bL32m in *GAL-MRX9* isolated mitochondria (Figure 3A). The processing of the mitochondrial presequence of bL32m is a crucial step required for the biogenesis of functional mitoribosomes and is orchestrated by the m-AAA proteases Yta10 and Yta12 [42]. Both Yta10 and Yta12 are essential components of the PHB/m-AAA complex [43], and mutations in their proteolytic centers or their deletion lead to an accumulation of immature bL32m, resulting in overall impairment of mitochondrial protein synthesis [42]. Previous studies have shown that strains harboring mutations in Yta10/Yta12 together with a bL32m variant whose processing is independent of m-AAA proteases retain respiratory competence and mitoribosome assembly, demonstrating the key role of these factors in bL32m maturation [42,44]. Notably, m-AAA proteases have been implicated in *COX1* and *COB* mRNA splicing [45]. To further examine the accumulation of the bL32m, we analyzed the steady-state levels of bL32m in isolated mitochondria from *GAL-MRX9*, Δ*yta10* and Δ*yta12* strains. Indeed, an unprocessed version of bL32m accumulates under high Mrx9 levels; however, its mass appeared to be less than observed in both protease mutants (Figure 3C). Nevertheless, this smaller precursor represents an immature form of bL32m since it was not detected in wild-type and Δ*bL32m* mitochondria and could potentially be a partially processed protein due to inefficient proteolysis; that is, excess Mrx9 did not completely abolish protease function (as in the *yta10 yta12* null mutants) but impaired it to some degree, leading to the accumulation of this aberrant bL32.

Altogether, our data indicate that although mitochondrial translation is impaired in *GAL-MRX9* mitochondria, mitoribosome assembly is upregulated as a compensatory response to a decreased rate of protein synthesis. Therefore, the observed defects in bL32m maturation and mitochondrial mRNA splicing point to a possible negative impact of Mrx9 excess on m-AAA protease function.

### Mrx9 interacts with the conserved prohibitin/m-AAA protease supercomplex at the polypetide exit tunnel of the mitoribosome

Our data suggest that Mrx9 behaves as a negative regulator of mitochondrial gene expression. To gain insights into the mechanism, we explored Mrx9 interactors by performing immunoprecipitation assays followed by proteomic analysis. To this end, we integrated a double-tagged version of Mrx9 (protein C peptide plus polyhistidine tags: “CH”) in the Δ*mrx9* mutant cells, in which expression is driven by the wildtype *MRX9* promoter. The recombinant protein Mrx9-CH, expressed at endogenous levels, was pulled down from mitochondrial extracts, and the eluate fraction was analyzed by mass spectrometry. Gene ontology analysis revealed that Mrx9 strongly associated with mitoribosomes and PHB/m-AAA complexes (Figure 4A). The PHB/m-AAA complex is located in the IMM, where it is strategically bound to the mitoribosome PET. It is formed by multiple copies of the Phb1 and Phb2 proteins, as well as the hexameric m-AAA proteases Yta10 and Yta12 [43]. Together, these components promote the correct folding of nascent polypeptides at the PET, or ensure their proteolysis if folding fails [28]. Previous studies using proximity-dependent biotin identification (BioID) reported the presence of Mrx9 in the vicinity of the PHB/m-AAA complex, however no direct interaction was demonstrated [28, 29]. To validate our findings, we integrated a functional HA-tagged version of prohibitin Phb1 in the Δ*phb1*Δ*mrx9/MRX9*-CH strain. Pull down of Mrx9-CH from mitochondrial extracts from this strain confirmed the presence of Phb1-HA in the bound fraction (Figure 4B), indicating that Mrx9 is physically interacting with the PHB/m-AAA complex components.

**Fig. 4.**
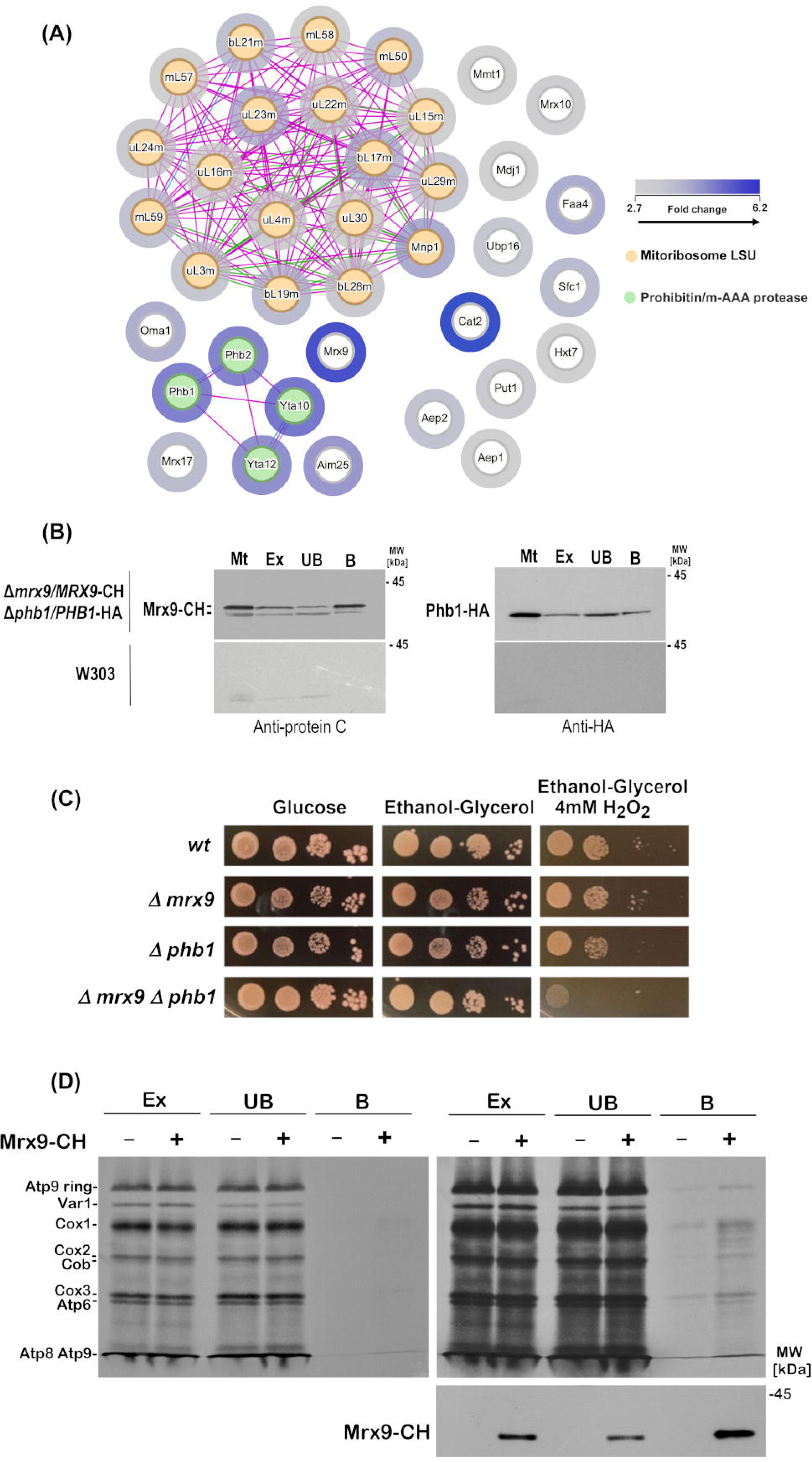
Mrx9 interacts with PHB/m-AAA complex factors, mitoribosomal proteins, and newly synthesized mitochondrial polypeptides. (A) Co-immunoprecipitation followed by mass spectrometry was performed using eluates from wild-type and Mrx9-CH pull down samples. A gene ontology interaction map of Mrx9 interactors was generated using the *String* platform, based on fold-change values. Pink lines mean experimentally determined interactions; blue lines represent known interactions from curated databases; green lines mean predicted interactions based on gene neighborhood; purple lines refer to protein homology. (B) Pull down assays using protein C–conjugated beads were performed with isolated mitochondria from wild-type and Δ*mrx9/MRX9*-CHΔ*phb1/phb1*-HA strains. Mitochondria (Mt), digitonin-extracted input (Ex), unbound (UB), and bound (B) fractions were separated by 12% SDS-PAGE. Anti-HA and anti–protein C antibodies were used for detection. (C) Growth properties of wild-type, Δ*mrx9*, Δ*phb1* and Δ*mrx9*Δ*phb1* strains at 30°C. Cells were spotted on rich glucose and rich non-fermentable (ethanol–glycerol) media containing or not 4 mM H O. Plates were photographed after 2 days of growth. (D) Co-immunoprecipitation of Mrx9-CH with newly synthesized mitochondrial polypeptides. Wild-type and Δ*mrx9/MRX9-*CH mitochondria were labeled with [³ S]-methionine/cysteine for 30 minutes at 30°C, followed by pull-down using protein C-conjugated beads. Digitonin-extracted input (Ex), unbound (UB) and bound (B) fractions were separated by 17.5% SDS-PAGE. The film on the left represents a shorter exposure, whereas the film on the right represents a longer exposure. The anti-protein C antibody was used to detect Mrx9-CH in the indicated fractions.

Based on its interactors, we hypothesized that *MRX9* may genetically interact with them, thereby reinforcing their functional association. Consistent with this, the *mrx9 phb1* double mutant exhibited a growth deficiency on ethanol–glycerol medium containing 4 mM H O, whereas neither single mutant showed any growth impairment under the same conditions (Figure 4C). This synthetic phenotype suggests that the combined loss of Mrx9 and Phb1 compromises cellular fitness under oxidative stress. One possible mechanism is a defect in mitochondrial proteostasis, for instance, in the absence of both proteins oxidatively damaged proteins may fail to be turned over properly.

In agreement with previous studies [28, 29], in our mass-spectrometry analysis we also detected accessory proteins of the PHB/m-AAA complex interacting with Mrx9. Among these factors are Oma1, a protease involved in the regulation of cellular stress responses and signaling [46], and two uncharacterized proteins, Mrx17 and Aim25 (Figure 4A). Interestingly, Oma1 is responsible for the degradation of newly synthesized Cox1 in cells with impaired Complex IV assembly, specifically due to the lack of the Coa2 assembly factor [47]. Consistent with the connection between Mrx9 and the quality control machinery, we also observed mitoribosome PET components in the Mrx9 interactome, including uL22m, uL23m and uL24m (Figure 4A). Additional putative Mrx9 interactors detected in our study are indicated in Figure 4A, including the highly enriched carnitine acetyl-CoA transferase Cat2. However, none of these interactors appeared to be associated with Mrx9 partners.

Because Mrx9 was found to interact with the PHB/m-AAA complex and proteins of the mitoribosome PET, we next asked whether it could interact with nascent mitochondrial polypeptides at the exit tunnel. To corroborate this hypothesis, translation products were labeled with ³ S-methionine/cysteine in mitochondria isolated from the Δ*mrx9/MRX9*-CH strain, followed by pull down analysis of the extracts. In the Mrx9-CH bound fraction, we observed an enrichment of newly translated polypeptides compared to the negative control (Figure 4D). Given that components of the PHB/m-AAA complex co-eluted with Mrx9 (Figure 4A), the observed association with newly translated polypeptides may be mediated indirectly through this complex. On the other hand, it remains possible that the Mrx9 factor itself interacts with nascent mitochondrial products. Taken together, these results and previous data [28, 29] show a close relationship between Mrx9, the PHB/m-AAA complex, and the mitoribosome. This connection suggests a role for Mrx9 in translational regulation and quality control.

### *MRX9* overexpression inhibits degradation of mitochondrial aberrant products

Considering that Yta10 and Yta12 are required for intron/maturase processing, which is essential for the splicing of *COX1* and *COB* introns [45], we hypothesized that the intron processing defects causing the appearance of the aberrant Cox1 polypeptide Mp15 upon *MRX9* overexpression might stem from impaired Yta10/Yta12 protease activity. Importantly, mitochondrial translation products were poorly detectable in the *yta10* null mutant, a phenotype most likely attributable to the lack of mature bL32m, as this ribosomal protein requires m-AAA protease activity for its proper processing (Figure S3A). Therefore, the perturbation in transcript splicing and bL32m presequence processing observed in cells overexpressing *MRX9*, together with the physical interaction between Mrx9 and Yta10/Yta12 detected by mass spectrometry, led us to hypothesize that Mrx9 may negatively modulate the functions of these proteases

On the other hand, previous studies showed that the m-AAA proteases are regulated by Phb1 and Phb2. Cells lacking prohibitins display accelerated degradation of non-assembled polypeptides by Yta10/Yta12, indicating that Phb1 and Phb2 might stabilize the m-AAA complex in an arrangement with reduced proteolytic activity [43].

To further investigate a possible role of Mrx9 in regulating m-AAA protease actions, similar to or in concert with prohibitins, we analyzed the stability of aberrant products that are substrates of Yta10/Yta12 in cells overexpressing *MRX9*. One suitable approach was to examine the stability of Mp15. This aberrant polypeptide has a stability of approximately 1 hour in *in vivo* pulse-chase experiments of mitochondrial translation, and its degradation is mediated by m-AAA proteases, as shown in previous studies [32]. We also took advantage of the fact that Mp15 accumulates not only under *MRX9* overexpression conditions but also in cytochrome *c* oxidase mutants such as cells lacking the *COX1* mRNA translational activator Mss51 [32,48, 49]. As a positive control, we tested the turnover of mitochondrial products and Mp15 in Δ*mss51* mutant cells. Indeed, Mp15 undergoes clear proteolysis after 1 hour of chase (Figure 5A). Confirming our hypothesis, cells overexpressing *MRX9* display strong resistance to Mp15 degradation even after 4 hours of chase (Figure 5B), suggesting an impairment in the function of the Yta10/Yta12 proteases. As a control we also measured Var1 stability over time.

**Fig. 5.**
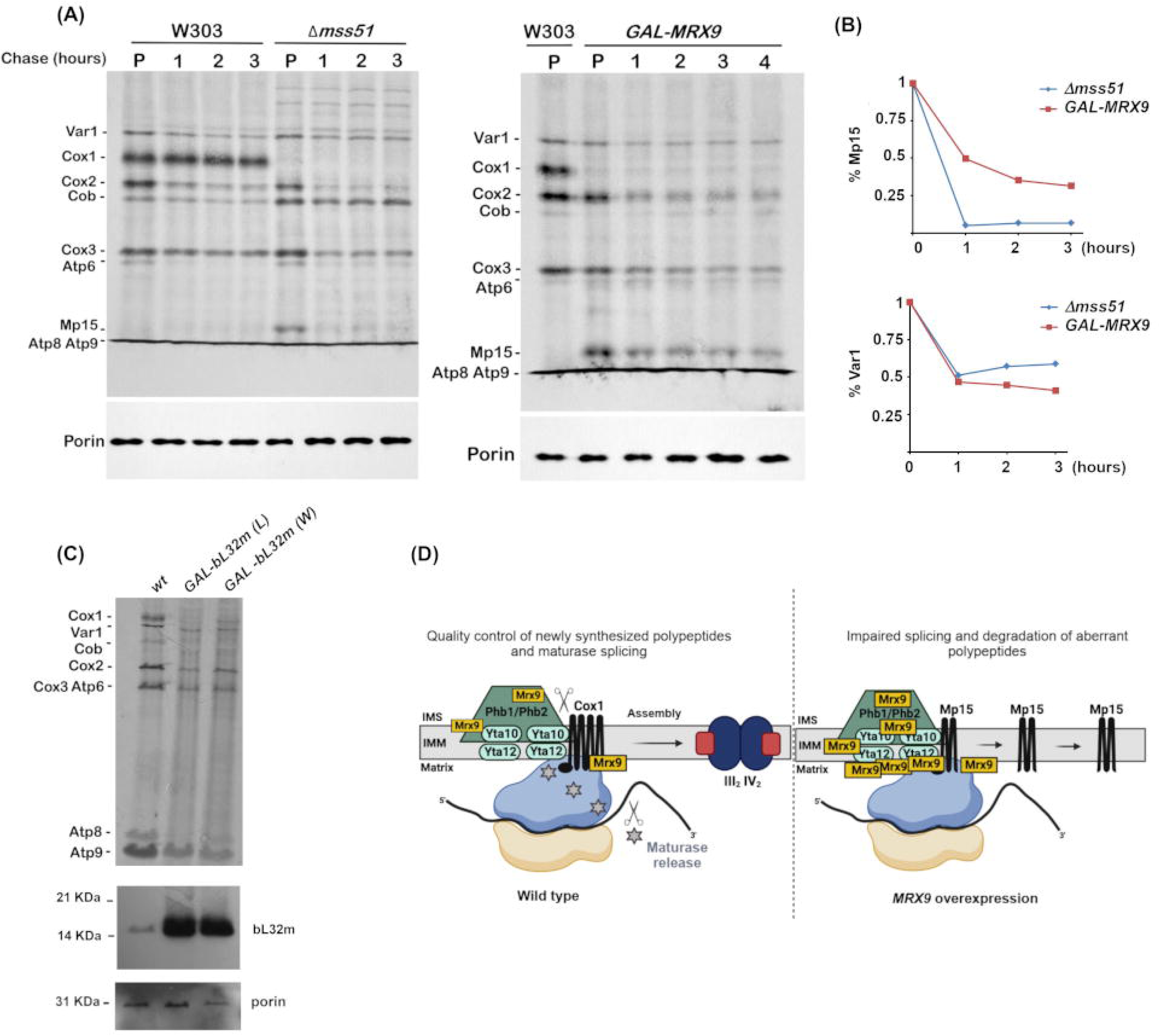
*MRX9* overexpression impairs Yta10/Yta12 proteolytic activity. (A, B) Analysis of newly synthesized mitochondrial translation products in W303, Δ*mss51* and *GAL-MRX9* cells. The indicated strains were pulse-labeled with [³ S]-methionine/cysteine for 20 minutes at room temperature. Labeling was stopped by adding 1000 μg of cold methionine/cysteine and 40 μg of puromycin. The stability of the translated polypeptides was assayed in chases of 1, 2, 3 and 4 hours at 30°C. Samples were separated by 17.5% SDS-PAGE. Porin was used as a loading control. (C) Analysis of newly synthesized mitochondrial translation products in W303, GAL-bL32m (L) and GAL-bL32m (W) cells. Overexpression of bL32m was analyzed using integrative plasmids with different selection markers. The indicated strains were pulse-labeled with [³ S]-methionine/cysteine as described in (A). Porin was used as a loading control. An antibody against bL32m was used to confirm the overexpression constructs. (D) Model of Mrx9 association with the PHB/m-AAA complex. In wild-type mitochondria, Mrx9 is located in the inner mitochondrial membrane, facing the matrix. Mrx9 is in proximity to the prohibitins Phb1 and Phb2, and two m-AAA proteases Yta10 and Yta12. The association of Mrx9 to the PET of the mitoribosome reflects its interaction with the PHB/m-AAA complex. Both m-AAA proteases are strategically positioned at the PET to facilitate the removal of introns and maturases from Cox1 and Cob nascent polypeptides, degrade aberrant or misfolded mitochondrial proteins and process the mitochondrial targeting sequence of the mtLSU protein bL32m. Excess Mrx9 appears to impair mitochondrial splicing and translation, with more pronounced effects on *COX1* and *COB* transcripts. Its overexpression leads to the stabilization of an aberrant truncated version of Cox1, named Mp15, which is not degraded by m-AAA factors. Overall, *MRX9* overexpression phenotype indicates a negative modulation of Yta10 and Yta12 proteolytic functions in mitochondria.

We also considered the possibility that Mrx9 may act as a natural substrate of the Yta10/Yta12 protease, and Mrx9 over accumulation could overload the protease, thereby impairing the degradation of other substrates such as the mitochondrial maturase. To test this hypothesis, we overexpressed bL32m, a well know substrate of these proteases. Therefore, bL32m was overexpressed from two different plasmids, and the accumulation of Mp15 was monitored by *in vivo* labeling of mitochondrial translation products (Figure 5C). However, overexpression of bL32m did not lead to any detectable increase in Mp15 levels and Cox1 itself was poorly labeled in these strain, indicating that excess bL32m causes a decrease in *COX1* translation that may precede Mp15 formation. We next asked whether the *yta10* null mutant alters Mrx9 steady-state levels. Yet, even when overexpressed, Mrx9 accumulated to similar levels in both wild-type and mutant cells (Figure S3B). These results suggest that excess Mrx9 likely exerts an inhibitory effect on m-AAA protease function rather than competing for substrate processing.

Furthermore, in yeast overexpressing *MRX9*, elevating *YTA10* expression alone did not alter Mp15 accumulation, suggesting that both Yta10 and Yta12 must be overexpressed in this context to mitigate the toxic effects of Mrx9p excess (Figure S4).

Mitochondrial translational fidelity and proteostasis have been linked to yeast lifespan [50]. We therefore sought to determine whether the physical association of Mrx9 with factors involved in mitochondrial protein quality control and translation confers a functional effect on yeast longevity. Analysis of *mrx9* mutants revealed no significant deviation from wild-type levels in either mean or maximal chronological lifespan, nor in viability assessed via the *in situ* survival assay (Figure S5) The same result was observed for *phb1* mutants in a previous study [51].

## Discussion

Mitoribosomes are responsible for translating the core subunits of the OXPHOS system complexes. Because mitochondrial polypeptides are highly hydrophobic and must be rapidly inserted into the IMM during biogenesis, these ribosomes are attached to the inner membrane in a sophisticated arrangement that couples translation and membrane insertion. Additionally, mitoribosomes are found in large assemblies associated with factors essential for mitochondrial gene expression, ranging from RNA processing and metabolism to translation activators and insertases. In *S. cerevisiae*, these mitoribosome assemblies were named MIOREX and are thought to connect transcript maturation and translation [3]. Interestingly, several studies in mammalian cells have demonstrated a similar organization of mitoribosomes, which are located near RNA granules containing posttranscriptional, translational and mitoribosome assembly factors [52–55].

In yeast, MIOREX complexes also include uncharacterized proteins whose functions remain to be elucidated [3]. Here, we unveil a regulatory role for the MIOREX protein Mrx9 in mitochondrial proteostasis and quality control through the modulation of m-AAA protease activity. In previous studies, we have linked Mrx9 to mitochondrial gene expression. Indeed, we have shown that while the absence of Mrx9 did not impair mitochondrial respiration, its overexpression resulted in impaired processing of the *COX1* and *COB* mRNAs [9]. Our current data further quantified the effects of Mrx9 on mitochondrial translation, revealing a general translation defect in cells overexpressing this protein, and suggesting that Mrx9 may act as a negative regulator of mitochondrial protein synthesis.

Translational repressors in mitochondria are poorly characterized, as their deletion mutants often show no detectable phenotype. One interesting example is Smt1, which functions as a translational repressor of the *ATP6/8* bicistronic mRNA [6]. By combining overexpression experiments with the analyses of mitochondrial protein synthesis, we have observed the disappearance of both Atp6/Atp8 translation in *GAL-SMT1* cells [9]. Here, we have followed a similar strategy to gain insights into the role of Mrx9 in mitochondrial gene expression. The data presented in this study identify three phenotypic consequences of high Mrx9 levels in living cells: a general translation impairment; a defect in *COX1* and *COB* mRNAs splicing; and an increase in the mitoribosome biogenesis, likely a compensatory mechanism to improve mitochondrial translation efficiency under stress caused by *MRX9* overexpression. Worth noting, Mrx9 overexpression does not affect the translation of Var1, a mitochondrion-encoded soluble SSU protein, the biogenesis of which does not require insertion in the IMM.

Based on our pull-down experiments, Mrx9 primarily binds to the mitoribosome LSU and the PHB/m-AAA complex, both large multi-protein structures. In the MitCOM complexome viewer [56], Mrx9 displays a distribution peak at approximately 2.2 MDa, and its overall distribution strongly correlates with that of the PHB/m-AAA proteins Phb1, Phb2, Yta10 and Yta12, strengthening the hypothesis that these factors coexist within the same complex. The PHB/m-AAA complex represents a point of integration between mitochondrial protein biogenesis and quality control. It forms a large ring-like structure [57], located in proximity to the mitoribosome PET [28,29], which acts as a docking platform preventing protein aggregation, misfolding, and proteotoxic stress during the translation of nascent polypeptides destined for insertion into the IMM [58]. Furthermore, the Yta10/Yta12 proteases are critical for the maturation of the LSU protein bL32m [41], and we reported here that Mrx9 excess results in a partial inhibition of bL32m processing. The presence of proteases near the PET is also a key feature that enables efficient excision of maturases encoded by mitochondrial introns. Interestingly, cells lacking both *PHB1* and *PHB2* genes are able to grow on non-fermentable carbon sources, whereas Δ*yta10* and Δ*yta12* mutants are respiratory deficient due to impaired mitoribosome biogenesis, resulting from defective bL32m processing, as well as impaired degradation of non-assembled inner membrane proteins and defective splicing [40,41,59]. Here, we observed a synthetic phenotype in the *phb1 mrx9* double mutant, reinforcing their putative functional association. To date, this is the most pronounced phenotype we have observed in *mrx9* deletion cells. While loss of prohibitins does not hypersensitize cells to endogenously produced ROS [51], the *mrx9 phb1* double mutant was sensitive to exogenous H O (Figure 4C), suggesting a role for these proteins in coping with external oxidative stress.

Moreover, previous proteomics studies indicated an approximately 75% decrease in Mrx9 abundance in Δ*yta10* and Δ*yta12* mutants compared to the wild-type strain, and a moderately strong positive correlation between the expression levels of Mrx9 and Yta12, with a Spearman’s rank correlation coefficient of 0.58 [55,60]. As for the PHB/m-AAA complex components, Mrx9 also might bind to newly synthesized polypeptides, associating Mrx9 roles with the quality control machinery of mitochondrial translation.

In summary, our results, together with multiple high-throughput studies [28,29,56,60], strongly suggest an association between Mrx9 and the PHB/m-AAA complex, with Mrx9 potentially behaving as a negative regulator of m-AAA protease activity. (Figure 5C). How Mrx9 acts precisely within this regulatory model remains unclear and warrant further studies.

## Materials and methods

### Yeast strains and growth media

All *S. cerevisiae* strains used in this study are listed in Table 1. For yeast culture media, cells were maintained in YPD (1% yeast extract, 2% peptone, 2% glucose), YPGal (1% yeast extract, 2% peptone, 2% galactose), YP-EG (1% yeast extract, 2% peptone, 2% glycerol, 2% ethanol), WO-Gal (2% galactose, 0.67% yeast nitrogen base), WO-Glu (2% glucose, 0.67% yeast nitrogen base). Strains were grown in liquid and solid media at 30°C.

**Table 1.**
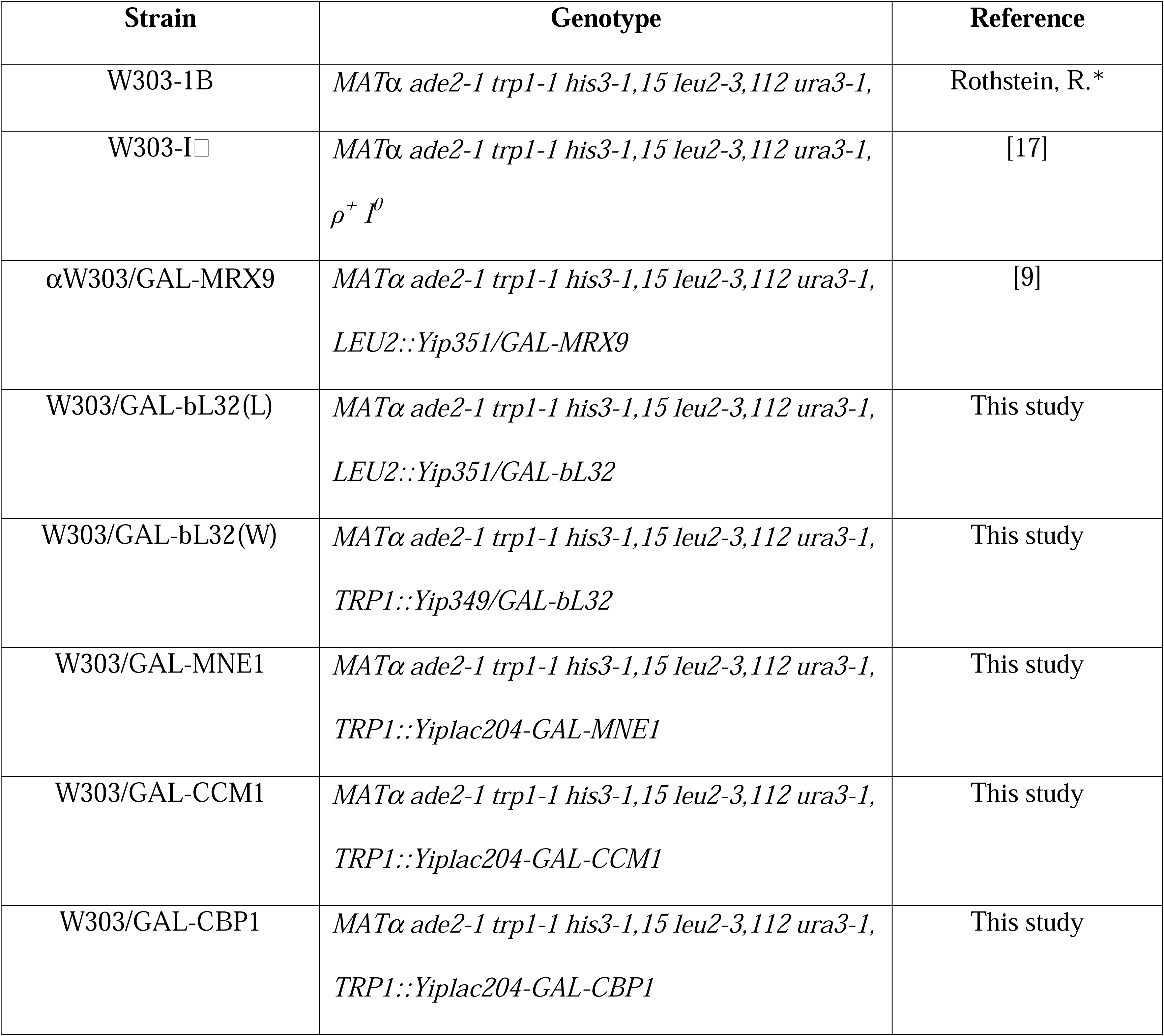
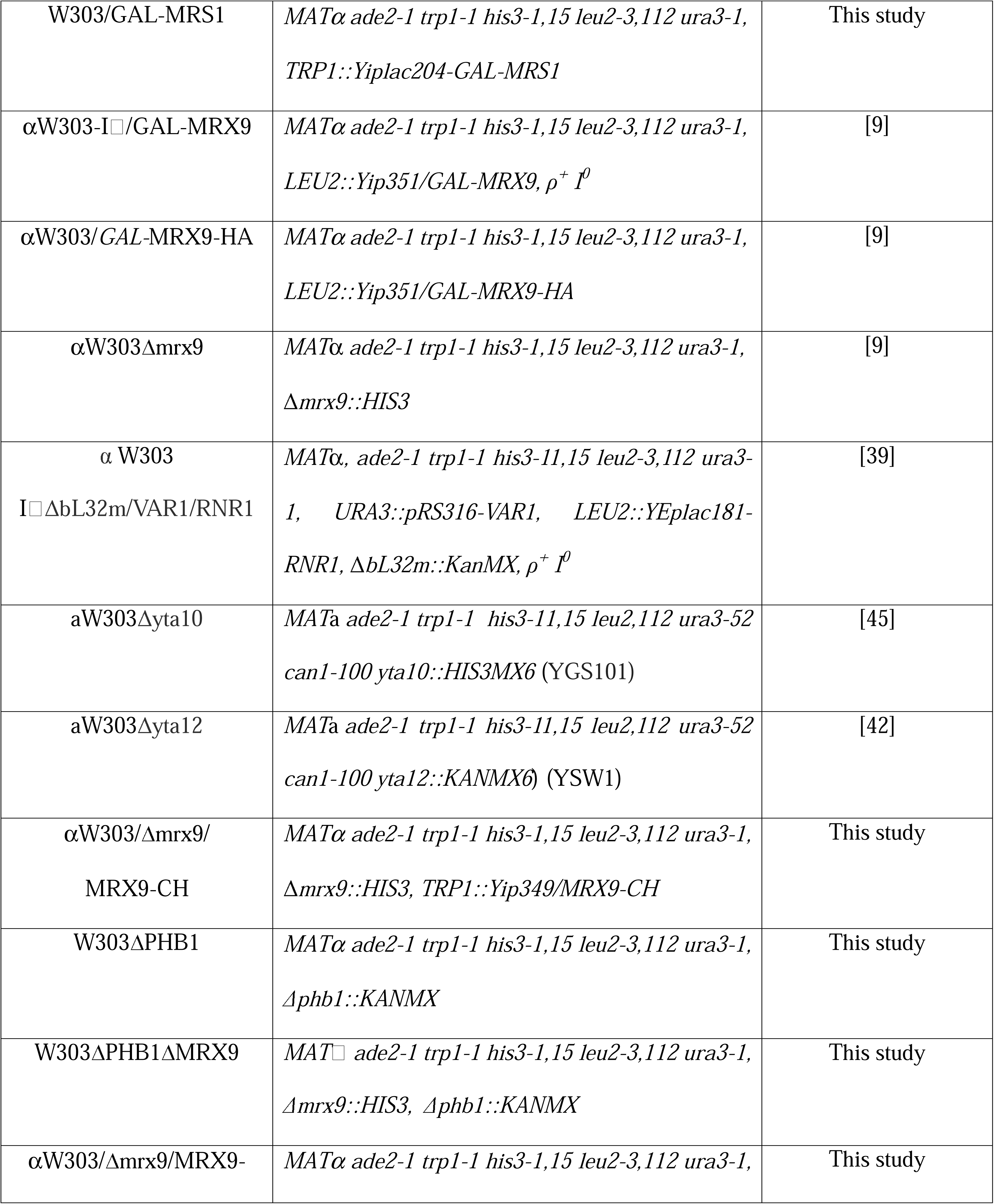
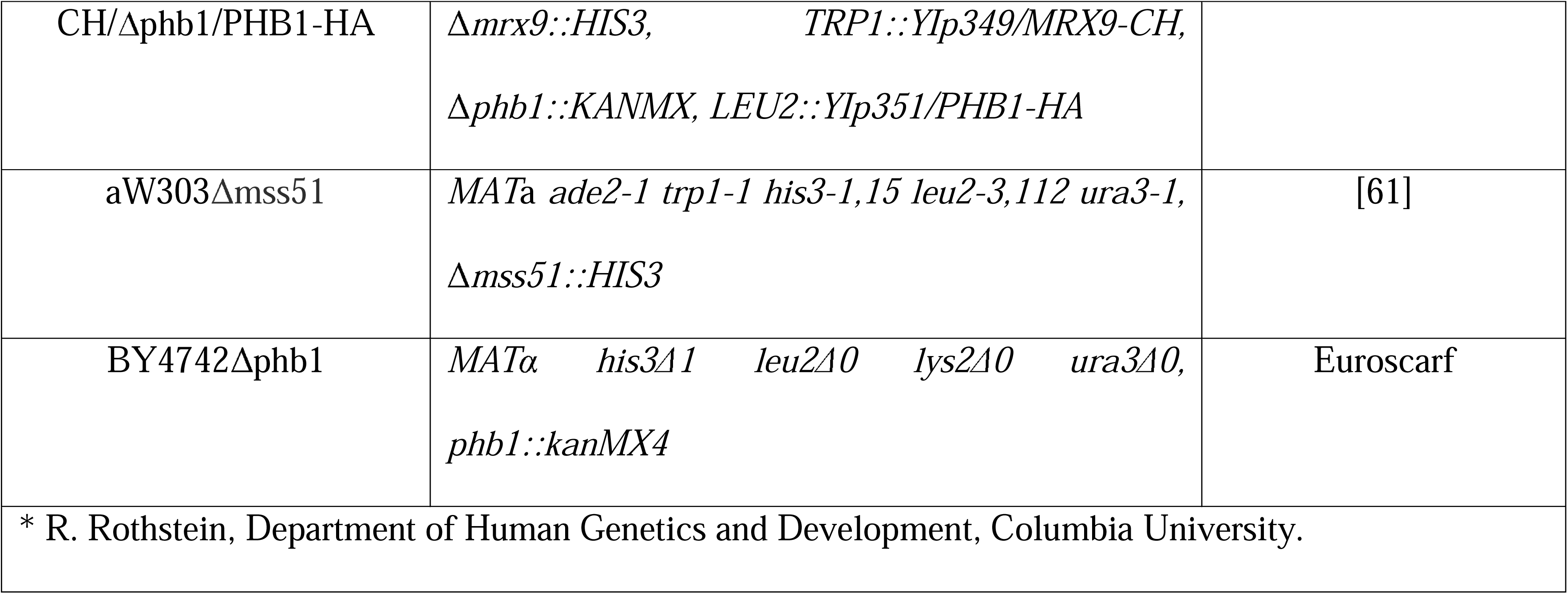
Yeast strains and relevant genotype used in this work.

### Construction of strains expressing Mrx9 double-tagged with protein C plus polyhistidine and HA-tagged Phb1

Mrx9 double-tagged with protein C followed by the polyhistidine tag was constructed by amplifying the gene with primers 5’-ggcagtcgacgaccaagcaacacgggaata-3’ and 5’-ggcgtcgacctcttttagggtgtattcag-3’. The PCR generated a product of approximately 1666 nucleotides, with 400 nucleotides of the 5’ untranslated region and the entire *MRX9* sequence minus the stop codon. The PCR product was digested with SalI and ligated into the plasmid YIp349-CH [62] cut with the same restriction enzyme. The *MRX9-CH* resulted in a frame fusion with the gene sequence and the CH tag. The epitope is composed of a Protein C tag followed by three glycines, six histidines and a stop codon. Phb1 expressing the hemagglutinin A (HA) tag was constructed by performing PCR using the primers 5’-ggcggatcccagtaagtggcagcataaccc-3’ and 5’-ggcctgcagtcaagcgtagtctgggacgtcgtatgggtaacggccaatgttcaaaag-3’. The PCR product of approximately 1260 nucleotides carries 400 nucleotides of the 5’ untranslated region, the entire *PHB1* minus the stop codon, and the HA tag sequence. The *PHB1*-HA fragment digested with BamHI and PstI was ligated into the plasmid YIp351 [63] cut with the same restriction enzymes. The recombinant plasmids *MRX9*-*CH*/YIp349 and *PHB1-HA*/YIp351 were linearized at their *TRP1* and *LEU2* markers, respectively, and integrated into the yeast chromosomal DNA by homologous recombination [64].

### Cloning and disruption of *PHB1*

The *phb1::KANMX4* cassette was generated by amplifying the fragment from the mutant BY4742Δ*phb1* obtained from Euroscarf. The PCR was performed using the primers 5’-ggcggatcccagtaagtggcagcataaccc-3’ and 5’-ggcctgcagccaggtttaagcctactgg-3’. The product was used to transform the W303Δ*mrx9/MRX9*-CH strain. To confirm the disruption, the DNA extracted from the strain was used in a PCR reaction with primers 5’-ggccgctttggctgtaggta-3’ and 5’-tgattttgatgacgagcgtaat-3’.

### Mitochondrial protein synthesis

Yeast cells were grown in minimum galactose media until reaching mid-log phase. Cytoplasmic translation was inhibited by adding 100 μg of cycloheximide. Mitochondrial gene products were labeled with [³ S]-methionine (7 mCi/mmol^−1^) or [³ S]-methionine/cysteine (7 mCi/mmol^−1^) at room temperature, as described previously [9,20]. Total cellular proteins with equivalent amounts were separated by 17.5% SDS-PAGE, followed by a nitrocellulose transfer and X-ray film exposure. *In organello* mitochondrial translation assay was performed using 2 mg of fresh isolated mitochondria. The translation buffer combined 0.6 M sorbitol, 150 mM KCl, 15 mM Kpi pH 7.4, 20 mM HEPES pH 7.4, 12.66 mM MgCl, 4 mM ATP, 0.5 mM GTP, 5 mM phosphoenolpyruvate, 5 mM α-ketoglutarate, amino acid mix (without Met, Tyr, Cys) with 12.13 μg/mL each amino acid, 12.13 μg/mL tyrosine and 10 μg/mL pyruvate kinase. Mitochondrial gene products were labeled with [³ S]-methionine/cysteine (7 mCi/mmol^−1^) at 30°C for 30 min, followed by the co-immunoprecipitation analysis [20,65].

### RNA isolation, RT-PCR and qPCR analysis

Total RNA was extracted from yeast cells with acidic phenol and SDS at 65°C, as described previously [66]. Before proceeding with RT-PCR assay, 10 µg of RNA were treated with 2 Units of DNase I/RNase-free (New England BioLabs) for 30 min at 37°C. 2 µg of RNAs were used for RT-PCR tests by using the High-Capacity cDNA Reverse Transcription Kit (Thermo Fisher Scientific). cDNA was quantified by qPCR using the universal SYBR Green fast qPCR mix (ABclonal) in a CFX96 Touch™ Real-Time PCR Detection System (Bio-Rad) as previously reported [20], adding primers described in Table 2. The experiment was performed in three independent repetitions.

**Table 2.**
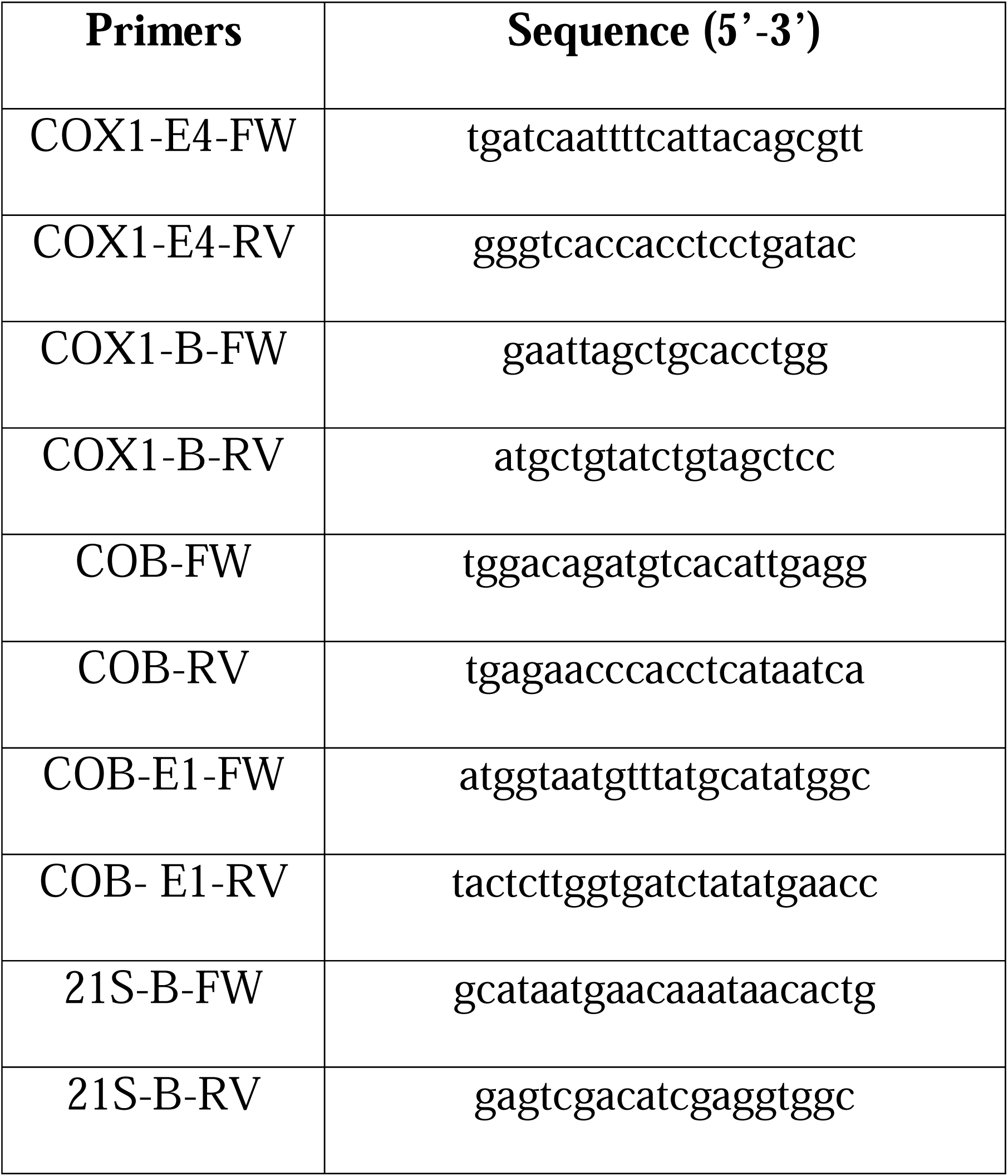
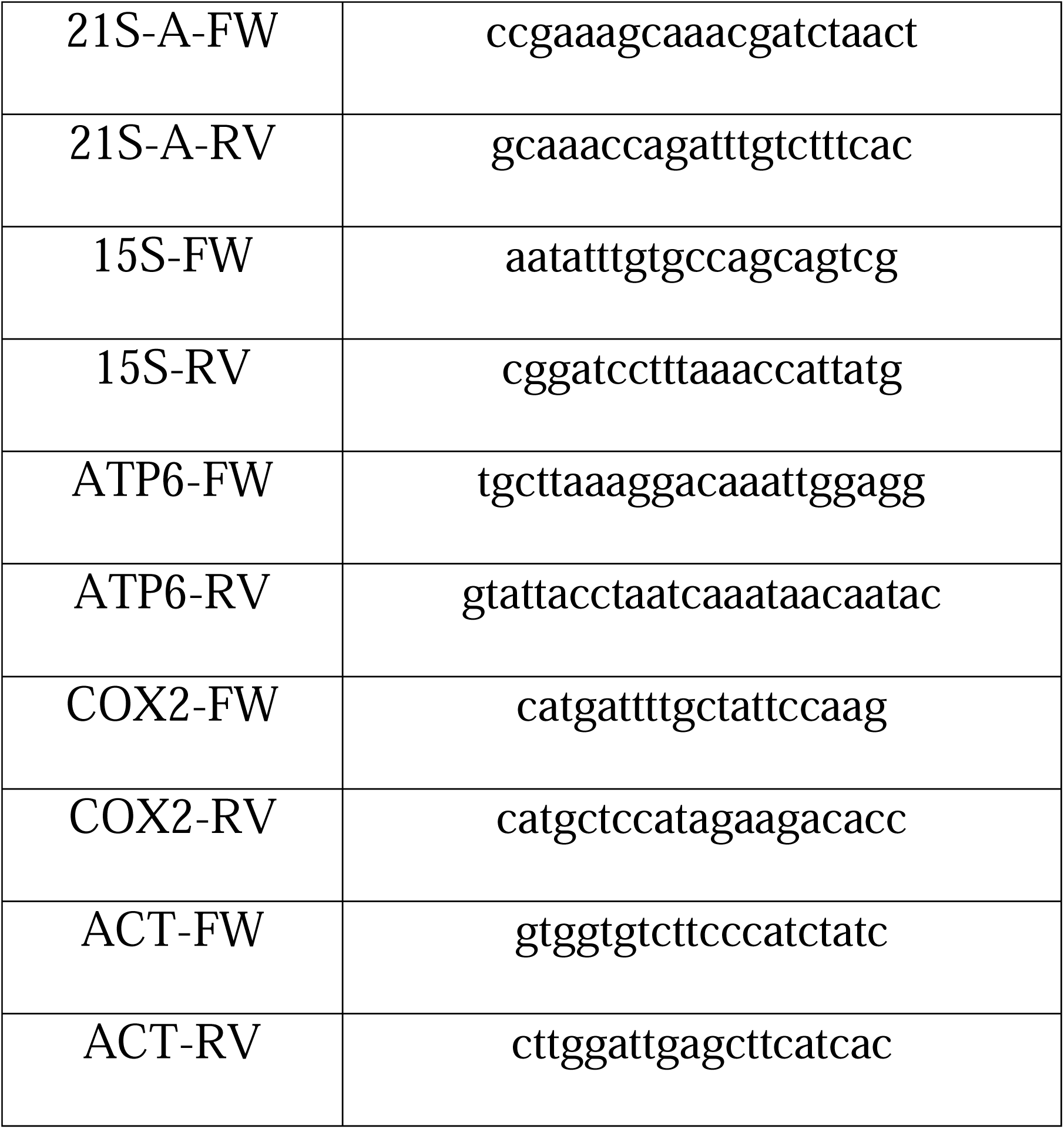
Primers used for qPCR analysis.

### Co-immunoprecipitation

Yeast mitochondria were isolated by the method of Herrmann et al. [67]. Mrx9-CH was purified by pull down experiments as described previously [6], with the following modifications. 2 mg of mitochondria were solubilized in 200 µl of extraction buffer (20 mM HEPES pH 7.4, 150 mM KAc, 2 mM 6-AHA, 0.5 mM PMSF, 0.8% digitonin) on ice for 5 min. The extract was obtained by centrifugation at 18,000 × g for 30 min at 4°C and incubated with anti-protein-C-conjugated beads (Roche) for 90 min at 4°C with gentle rocking. Supernatants (unbound fraction) were recovered after centrifugation at 500 × g for 5 min and beads were washed 4–5 times with wash buffer (20 mM Tris-Cl pH 7.5, 100 mM NaCl, 2 mM CaCl, 0.2% digitonin). The Mrx9-CH was eluted with elution buffer (20 mM Tris-Cl pH 7.5, 50 mM NaCl, 7 mM EDTA pH 7.5, 0.2% digitonin) for 30 min at 4°C with gentle rocking. The eluate was recovered after centrifugation at 500 × g for 5 min. Extract (input), unbound and bound samples were analyzed by Western blot.

### Mass spectrometry and bioinformatics analyses

Mass spectrometry analyses were performed by analyzing the eluate fraction from co-immunoprecipitation assays at the Redox Proteomics Core of the Mass Spectrometry Resource at the Chemistry Institute, University of São Paulo. Data were processed using the Perseus software suite (http://www.perseus-framework.org/). Following log transformation, proteins were filtered to require at least 50% valid values in either the *MRX9-CH* or wild-type group. A two-sample t-test was then employed to compare mean label-free quantitation (LFQ) intensities between the two groups.

### Life span and viability assays

Chronological life span (CLS) and in situ survival were measured as previously described [68]. Briefly, yeast cells were grown to stationary phase in liquid media, and viability was monitored periodically by determining the fraction of cells capable of forming colonies (CFU) on YPD plates. For each culture, Abs600 was measured, serial dilutions were prepared to a final Abs600 of 0.001, and 33 μL of this dilution was spread onto YPD plates. Plates were incubated for 72 h at 30 °C to allow colony formation, after which CFU were counted.

Survival was also assessed using a viability-on-a-plate *in situ* assay based on the leucine auxotrophy of the tested strains. Abs600 was determined for each culture, serial dilutions were prepared to a final Abs600 of 0.001, and 33 μL of this dilution was spread onto ten minimal media plates supplemented with all auxotrophic requirements except leucine. At time zero, one plate per strain was supplemented with tryptophan, and the total number of CFU was counted. All plates were incubated at 30 °C, and leucine was added to the remaining plates every one or two days to allow viable cells to grow.

In both assays, experiments were performed in triplicate, and viability was expressed as a percentage of the CFU count at time zero (defined as 100% viability).

### Miscellaneous procedures

Protein concentrations were quantified by the method of Lowry [69] and samples were suspended in Laemmli buffer [70] for SDS-PAGE analyses. Antibodies used for steady-state level experiments are listed in Table 3. Densitometry analysis of the bands was performed by using the ImageJ platform. GraphPad Prism software (GraphPad Software, USA) was used for graphs and data analyses. The Tukey test was performed for three groups and Student’s t-test for two groups. A p value above 0.05 was considered significant.

**Table 3.**
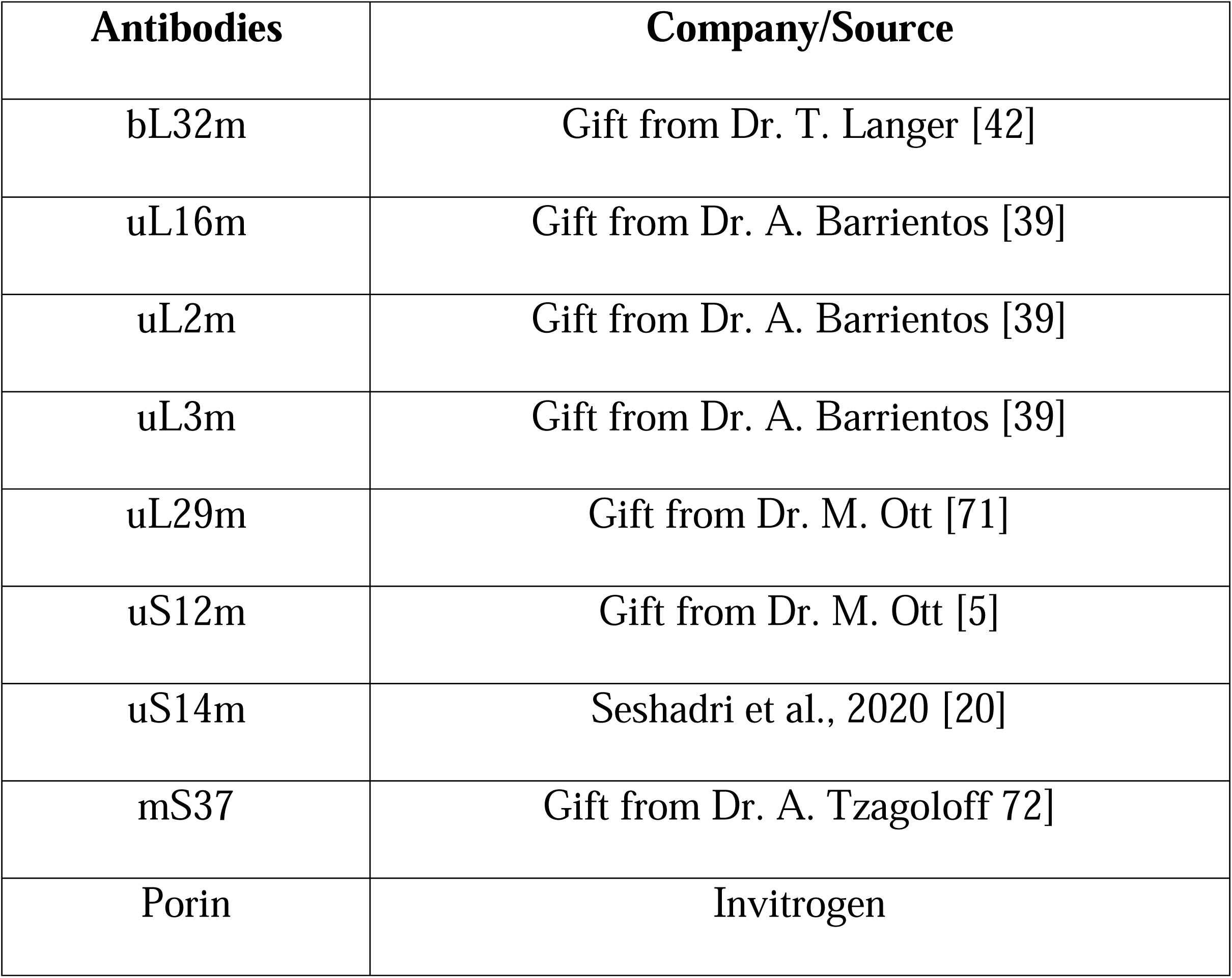
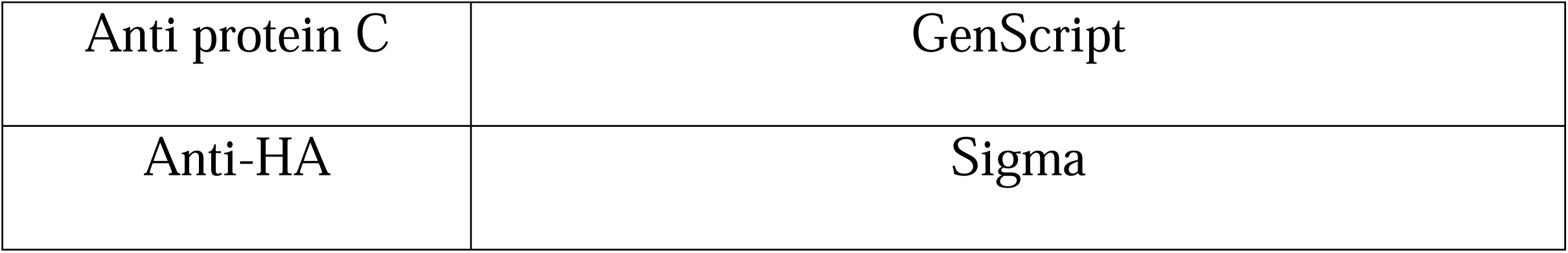
Antibodies used in this work.

## Acknowledgments

We thank Professors Paolo Di Mascio and Graziella E. Ronsein for the mass spectrometry analyses performed at the Redox Proteomics Core of the Mass Spectrometry Resource at Chemistry Institute, University of São Paulo (FAPESP grant numbers 2012/12663-1, 2016/00696-3, CEPID Redoxoma 2013/07937-8), and Dr. Mariana P. Massafera for her technical assistance. We also thank Dr Carol L. Dieckmann for critically reading this manuscript.

## Author Contributions

JACC – Conceived, designed and performed the experiments, analyzed the data, prepared the digital images, and drafted the manuscript. FF - Conceived and designed experiments, and drafted the manuscript. ACPA – Performed experiments. MHB – Conceived and designed experiments, and drafted the manuscript.

