## Supplementary material for "The mitoribosome-associated factor Mrx9 acts as a negative regulator of the prohibitin/m-AAA complex": Figure S1

**Figure S1** – *In vivo* labeling of mitochondrial products from cells grown in minimal galactose media. A) Comparison of the wild type strain (W303) and the transformant harboring YIp351-GAL plasmid (W303+ev). B) Analyses of *MRX9* overexpression (*GAL-MRX9*). together with factors involved in *COB* and *COX1* splicing – *GAL-MNE1*, *GAL-CCM1*, *GAL-CBP1* and *GAL-MRS1*. Mitochondrial products are indicated on the left of panel.

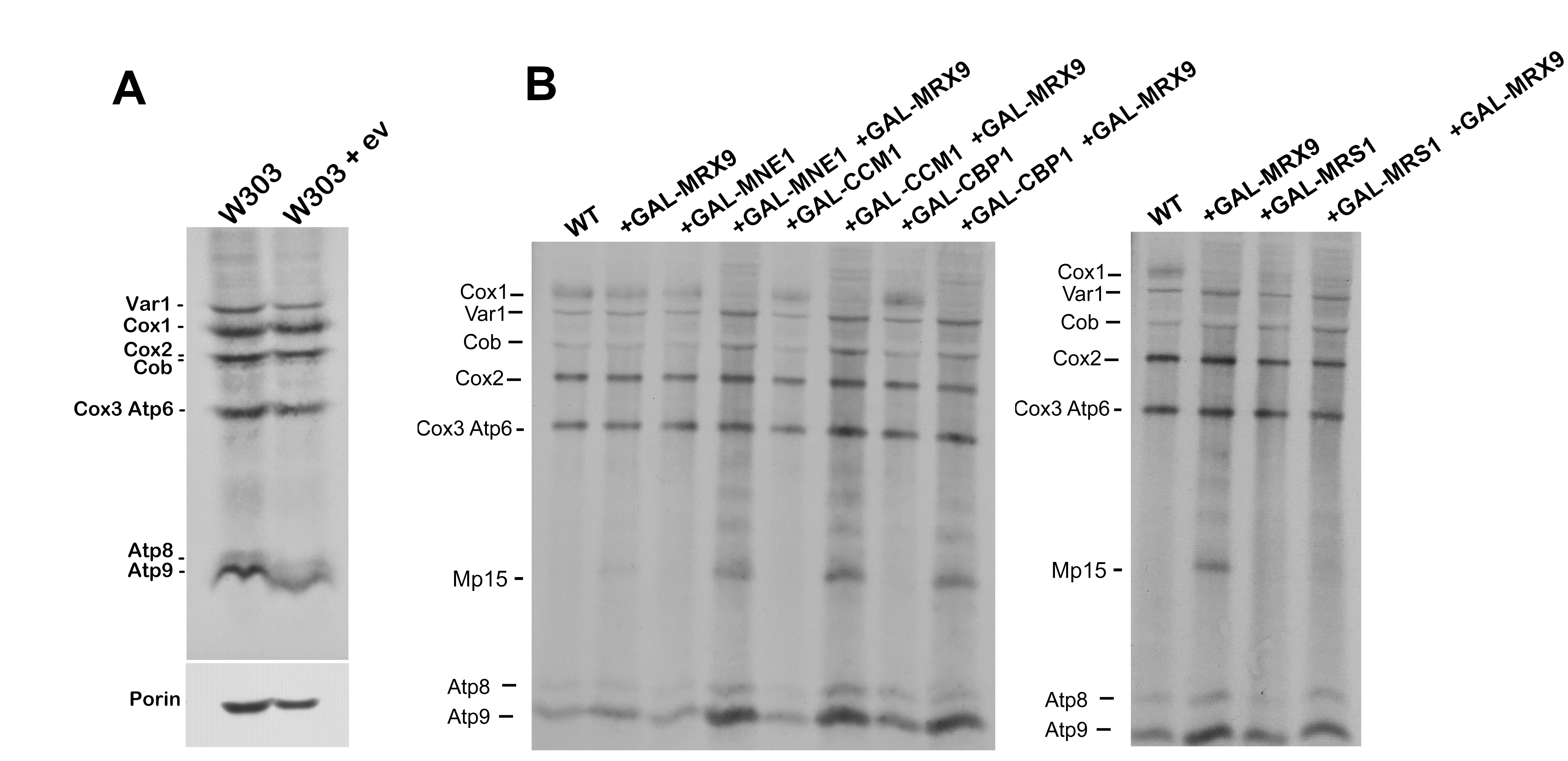

**Figure S2:** Three independent experiments of *in vivo* labelling of mitochondrial products of the following strains: the *mrx9* null mutant and the wild type (W303) containing or not introns (W303-I^o^) and transformed (+), or not (-) with the *GAL-MRX9* construct. Mitochondrial products are indicated on the left of panel. Band intensities of Cox1, Cox2, Cob, Cox3 and Atp6 were also compared using Var1 product to normalize the densitometry of the Adobe Photoshop program. The values obtained were compared in the bar graph shown on the bottom, * represents Student’s t-test statistics significance between the two compared values (p < 0.05).

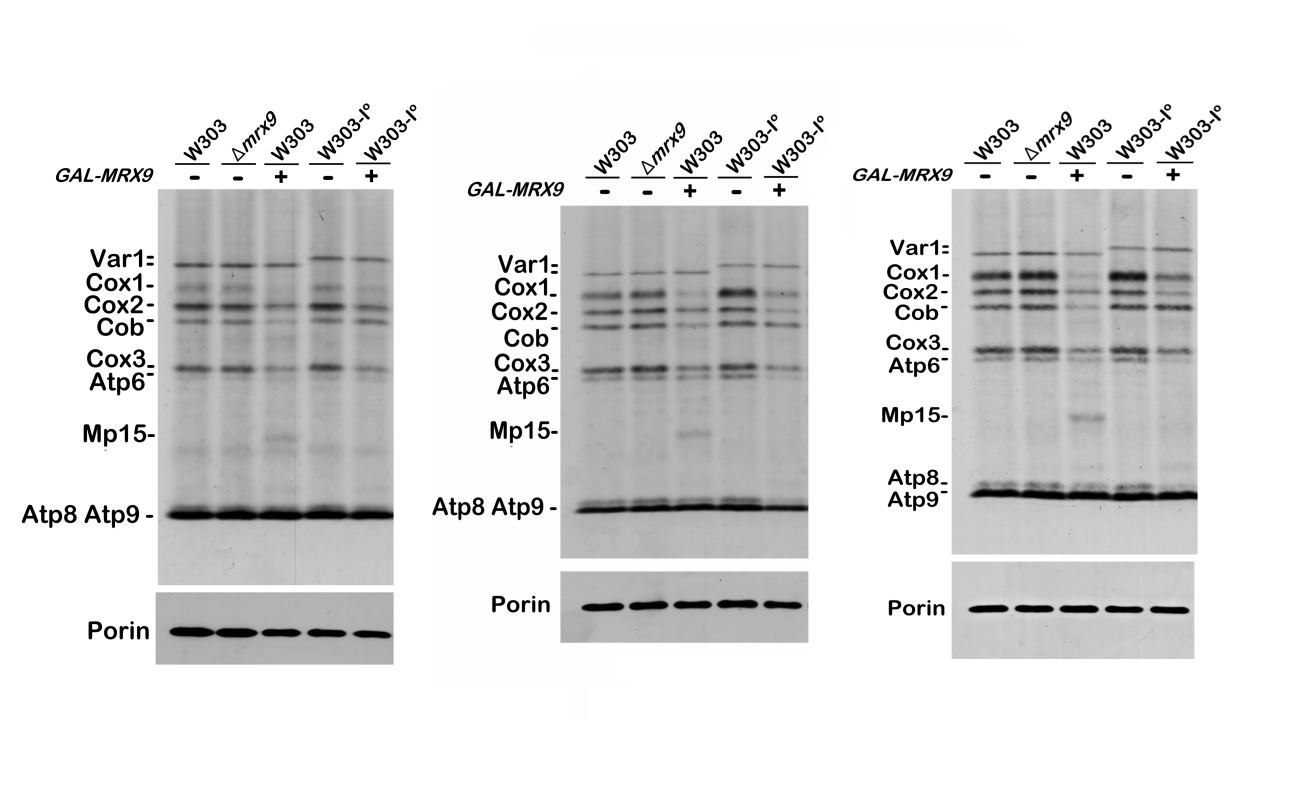

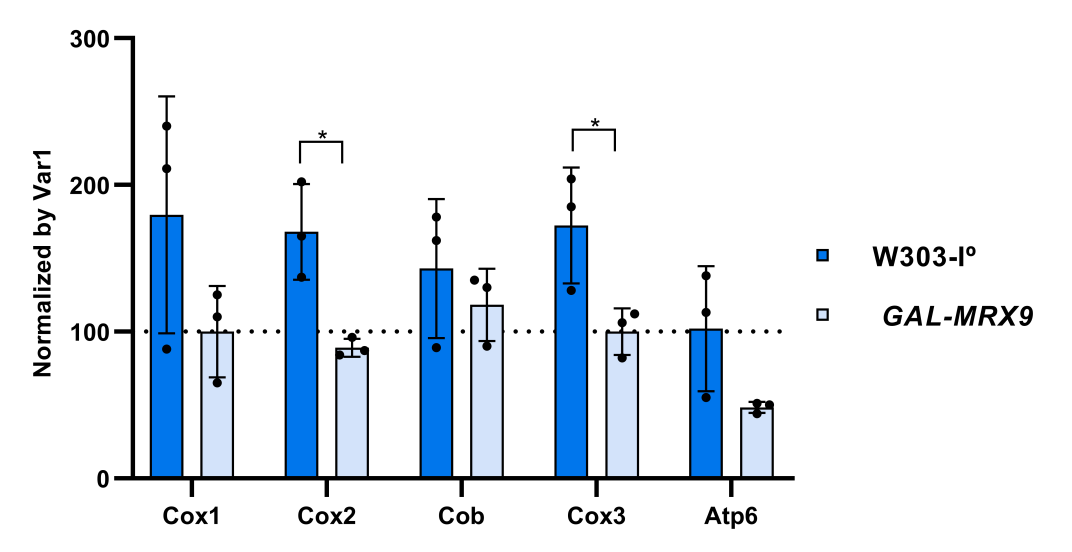

**Figure S3 – A)** *In vivo* labeling of mitochondrial products from cells grown in minimal galactose media. Mitochondrial products were labeled in the wild type strain (WT) the strain overexpressing *MRX9* (*GAL-MRX9*) and in the *yta10* null mutant (B) Mrx9p steady-state levels were assessed in wild-type and *yta10Δ* cells, either transformed or not with the *GAL-MRX9* overexpression construct. Molecular weight standards are indicated on the left of the respective panels.

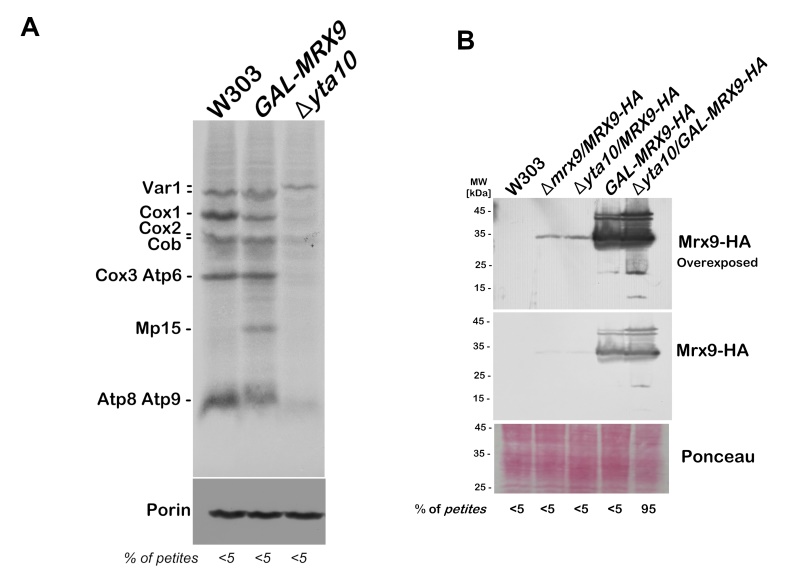

**Figure S4 -** *In vivo* labeling of mitochondrial products from cells grown in minimal galactose media and chased for four hours. A) Mitochondrial products were labeled in the wild type strain and in the strain over expressing *MRX9* and *YTA10* together. Labeling was stopped by adding 1000 μg of cold methionine/cysteine and 40 μg of puromycin. The stability of the translated polypeptides was assayed in chases of 2 and 4 hours at 30°C. Samples were separated by 17.5% SDS-PAGE. Porin was used as a loading control. B) Percentage of Mp15 and Var1 bands intensity in comparison to time 0 (pulse - P). Band densitometry counts were obtained using the histogram component of GNU Image Manipulation Program GIMP 2.10.28

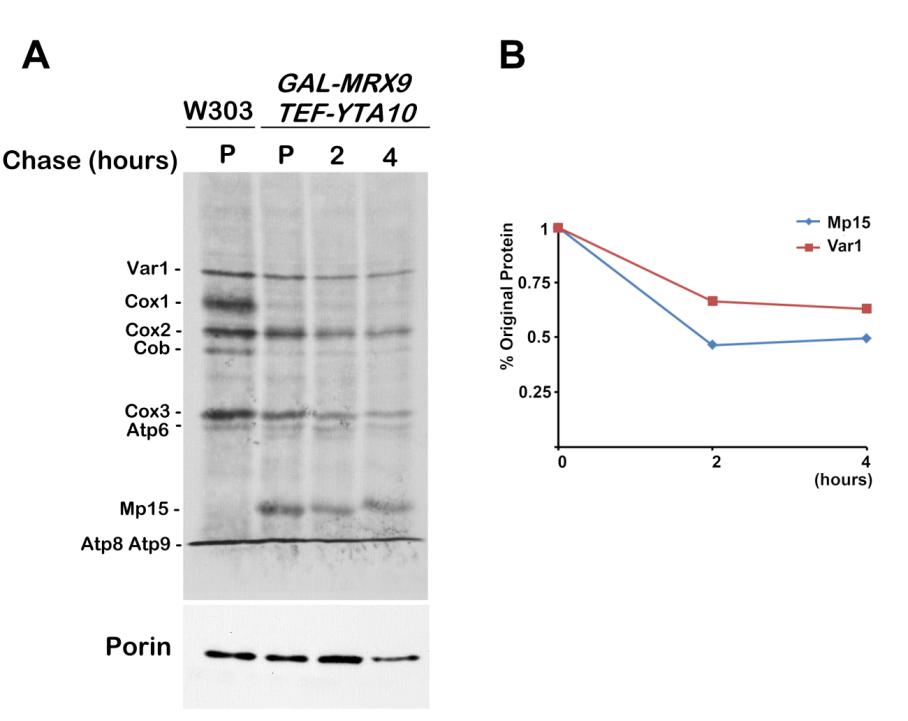

**Figure S5–** Viability assays of the wild type (wt) and *mrx9* null mutant. The results shown were calculated as the percentage of cell survival (number of colonies) relative to the counting of each strain at time zero, set as 100%. The results are expressed as the means +/- SD of three independent experiments run in duplicate. (A) Chronological Life Span. (B) *In situ* viability assay.

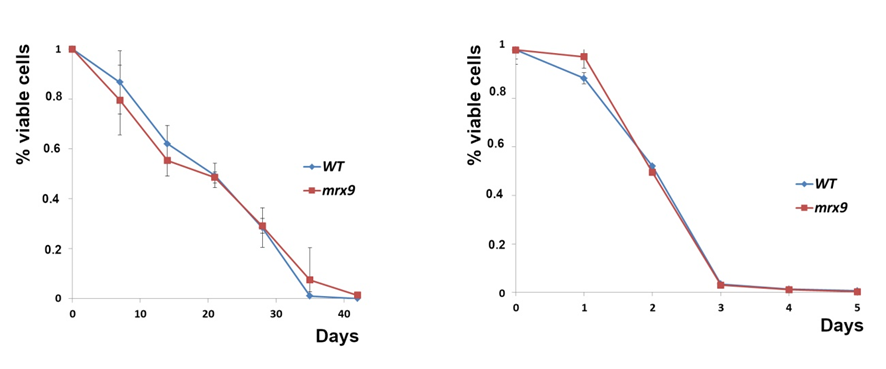
